# Rest functional brain maturation during the first year of life

**DOI:** 10.1101/760843

**Authors:** Hervé Lemaître, Pierre Augé, Ana Saitovitch, Alice Vinçon-Leite, Jean-Marc Tacchella, Ludovic Fillon, Raphael Calmon, Volodia Dangouloff-Ros, Raphaël Lévy, David Grévent, Francis Brunelle, Nathalie Boddaert, Monica Zilbovicius

## Abstract

The first year of life is a key period of brain development, characterized by dramatic structural and functional modifications. Here, we measured rest cerebral blood flow (CBF) modifications throughout babies’ first year of life using arterial spin labeling magnetic resonance imaging sequence in 52 infants, from 3 to 12 months of age. Overall, global rest CBF significantly increased during this age span. In addition, we found marked regional differences in local functional brain maturation. While primary sensorimotor cortices and insula showed early maturation, temporal and prefrontal region presented great rest CBF increase across the first year of life. Moreover, we highlighted a late and remarkably synchronous maturation of the prefrontal and posterior superior temporal cortices. These different patterns of regional cortical rest CBF modifications reflect a timetable of local functional brain maturation and are consistent with baby’s cognitive development within the first year of life.

## Introduction

The human brain is still immature at birth and undergoes dynamic structural and functional processes throughout life. During the first year, the maturation of neural networks is a complex process that is particularly important to the baby’s acquisition of cognitive and motor skills (Kagan and Herschkowitz 2005). At the cortical level, development comprises both gross morphometric changes and microstructural progression (Dubois et al. 2014). The first year of life is therefore a critical phase of postnatal brain development.

Historically, much of what we know about the intricate processes of early brain development comes from post-mortem studies in human fetuses, neonates, and non-human primates (Goldman-Rakic 1987; Kostovic et al. 2002; Innocenti and Price 2005). With the increasing availability of high-quality neuroimaging techniques, studying early human brain development in vivo in unprecedented detail is now feasible (Partridge et al. 2004; Fransson et al. 2007; Ball et al. 2014; van den Heuvel et al. 2015). These advances have led to exciting new insights into both healthy and atypical macroscale brain network development and have paved the way to bridge the gap between the brain’s neurobiological architecture and its behavioral repertoire.

At the structural level, in neonates and infants, studies of cortical morphological development have focused on the modification of gray matter volume (Knickmeyer et al. 2008; Gilmore et al. 2012), gyrification (Li et al. 2014), deep sulcal landmark maturation (Meng et al. 2014), thickness and surface area maturation (Lyall et al. 2015), as well as folding and fiber density (Nie et al. 2014). Structural brain imaging studies showed an increase in the gray matter volume during the first years of life (Knickmeyer et al. 2008), consistent with post-mortem studies, indicating rapid development of synapses and spines during this period (Huttenlocher and Dabholkar 1997; Petanjek et al. 2011; Webster et al. 2011). Indeed, throughout late gestation, rapid synaptogenesis results in an over-abundance of synapses (up to 150% of adult values) that are subsequently pruned throughout childhood and adolescence (Huttenlocher 1979). During the first year of life, synaptogenesis is one of the most important maturational processes, and its timetable differs across cortical regions. Gilmore et al. described a posterior to anterior gradient of gray matter growth throughout the first year of life (Gilmore et al. 2007), consistent with regional differences that have been described in post-mortem studies, showing an increase in synaptic density, and therefore synaptogenesis, earlier in the sensory cortex and later in the prefrontal cortex (Huttenlocher and Dabholkar 1997). In general, studies have suggested a complex pattern of development that varies based on anatomical location and cortical metrics. In addition, across early development, cortical maturation exhibits regionally specific asymmetry between the left and right hemispheres (Li et al. 2014; Nie et al. 2014). These changes continue throughout childhood and adolescence, with cortical thickness following different trajectories of thinning depending on the region, cortex type and gender (Sowell et al. 2004; Shaw et al. 2008).

At the functional level, early brain development has been investigated using mainly three different approaches. Pioneer studies measuring metabolism and rest cerebral blood flow (CBF) were followed by activation studies using functional MRI and more recently resting state MRI studies investigating functional connectivity.

Rest cerebral metabolism and blood flow are an index of synaptic density, which allows the in vivo study of functional brain maturation using positron emission tomography (PET) and single-photon emission computed tomography (SPECT) (Leenders et al. 1990). These studies revealed that infants’ brains showed higher rest metabolism in subcortical structures and in the sensorimotor cortex than in other regions (Chugani and Phelps 1986). In the newborn, the highest degree of glucose metabolism is in the primary sensory and motor cortex, cingulate cortex, thalamus, brain stem, cerebellar vermis and hippocampal region. During the first months of life, rest metabolism and CBF increase firstly within the primary sensory cortices, followed by the associative sensory cortices and finally within the prefrontal cortex at the end of the first year (Chugani and Phelps 1986; Chugani et al. 1987; Chiron et al. 1992). At 2 to 3 months of age, glucose utilization increases in the parietal, the temporal and the primary visual cortices, basal ganglia, and cerebellar hemispheres. Between 6 and 12 months of age, glucose utilization increases in the frontal cortex. These metabolic changes correspond to the emergence of motor and cognitive abilities during the first year of life. However, these studies were limited by very low spatial resolution of the brain imaging devices. In addition, these techniques required administration of ionizing radiation and, therefore, have limited application in the pediatric population.

Following these pioneer studies, functional MRI studies have used blood oxygen level-dependent (BOLD) contrast to measure brain activity. Task-based fMRI contributed to present-day knowledge about brain maturation shortly after birth (Dehaene-Lambertz et al. 2006; Arichi et al. 2010; Allievi et al. 2016). These studies have provided important background on the brain’s responses to sensory input during the early developmental phases of brain-behavior interactions. Adult-like activation patterns were observed in response to a variety of sensory stimuli, including tactile and proprioceptive stimulation (passive hand movement) (Erberich et al. 2006; Arichi et al. 2010) as well as auditory (Anderson et al. 2001) and olfactory (the odor of infant formula) (Arichi et al. 2013). Functional MRI studies in 2-to 3-month-old infants demonstrated left-lateralized activation of perisylvian regions, including the superior temporal gyrus, angular gyrus and Broca’s area, in response to native language speech. The response followed a hierarchical pattern, with auditory regions being activated first, followed by superior temporal regions, the temporal poles and Broca’s area in the inferior frontal cortex; a pattern that is highly consistent with language organization in the mature brain (Dehaene-Lambertz et al. 2006).

More recently, BOLD signal has been used to conduct resting state fMRI (rs-fMRI) as a measure of temporal coherence between brain regions. This technique has provided, by studying infants during the first years of life, insight into the maturation of multiple resting state networks (RSNs). Results show that the rate at which correlations within and between RSNs develop differs by network and closely reflect known rates of cortical development based on histological evidence (Smyser and Neil 2015). The sensorimotor (SM) and attention (AN) networks seem to be the earliest developing networks with their within-network synchronization largely established before birth. This replicates several reports showing the bilateral symmetric, adult-like topology of both networks at birth (Gao et al. 2013; Lin et al. 2013) or even prenatally (Smyser et al. 2010), indicating significant prenatal development of these 2 networks. In the brains of term babies, rs-fMRI studies employing seed-based connectivity or independent component analyses have identified specific functional networks, including primary visual, auditory, sensorimotor networks and default mode and executive-control networks involved in heteromodal functions (Fransson et al. 2007; Doria et al. 2010; Smyser et al. 2010). Network analyses based on graph theory further revealed that the functional connectomes of infant brains already exhibited the small-world structure. Distinct from the adults, however, the hubs were largely confined to primary sensorimotor regions (Fransson et al. 2011; Gao et al. 2011). Taken together, these findings provide important insights into the early brain functional maturation process.

The emergence of arterial spin labeling (ASL), a technique that provides both non-invasive and regional cerebral blood flow quantification, offers new opportunities to investigate local rest brain function in neonates and children. ASL perfusion MRI uses magnetically labeled arterial blood water as a nominally diffusible flow tracer. By labeling the blood water proximal to the target imaging region, the perfusion signal is subsequently calculated by comparison with a separate image acquired using a control pulse without labeling the blood flow to remove the static background signal and control for magnetization transfer effects (Williams et al. 1992). Therefore, ASL MRI non-invasively assesses brain perfusion and allows for a quantitative measurement of rest CBF without the administration of contrast material or exposure to ionizing radiation (Detre and Alsop 1999). Compared with BOLD, ASL has several benefits by providing (1) an absolute quantitative measure of brain function through CBF signaling, (2) not a derived measure from the BOLD signal affected by numerous physiological and noise contributions, and (3) an increased spatial specificity to neuronal activity due to the capillary/tissue origin of the ASL signal (Detre et al. 2009; Chen et al. 2015). Then, ASL MRI has been used to measure rest CBF to better characterize brain maturation in children after the first year of life (Biagi et al. 2007; Paniukov et al. 2020). Indeed, in a very recent study, Paniukov et al. (Paniukov et al. 2020) followed longitudinally children between 2 and 7 years and showed a constant CBF increase across different regions of the prefrontal, temporal, parietal and occipital cortex. However, knowledge about the first year of life remains rather scarce. Wang et al. (Wang et al. 2008) by comparing 7- and 13-month old infants showed a regional CBF increase in the hippocampi, anterior cingulate, amygdala, occipital lobe and auditory cortex. Duncan et al. (Duncan et al. 2014) studied infants from 3 to 5 months and described a significantly greater rest CBF in the orbitofrontal, subgenual and inferior occipital regions.

In this study, in order to characterize CBF developmental trajectories, we have measured age-related changes of local rest CBF at the voxel level and regionally throughout the first year of life using ASL perfusion MRI. We hypothesized that this crucial age range is characterized by different regional patterns of brain development, mainly between primary and associative regions, that in turn reflect cognitive development of the baby during the first year.

## Materials and Methods

### Subject

Eighty-five babies from the Necker-Enfants-Malades hospital were initially included in this study. The inclusion criteria were normal clinical multimodal MRI, absence of prematurity, neurological or cranial pathology, parent’s consanguinity or abnormal psychomotor development. Were included infants presenting syndromes that are not originally neurological, mainly dermatological or ophthalmological, but request an MRI to discard infrequent associated brain abnormalities, that may be present in a small percentage of cases (see SI Appendix, Table S1). Normal psychomotor development was assured in follow-up consultations. Our final sample included 52 babies (29 girls) from 3 to 12 months of age in our study, including 10 babies at 3 and 4 months (90 to 120 days), 14 at 5 and 6 months (120 to 180 days), 14 at 7 and 8 months (180 to 240 days), 7 at 9 and 10 months (240 to 300 days), 7 at 11 and 12 months (300 to 375 days). The Ethical Committee of French Public Hospitals approved this study and the written informed consent was obtained for all participants.

### MRI acquisition

All MRI exams included whole brain T1-weighted and ASL sequences and were acquired on a General Electric Signa 1.5T MRI scanner in the Necker-Enfants-Malades hospital (See SI Appendix, Table S2 and SI Methods for details). Due to the age of the babies, all of them received premedication before their MRI (pentobarbital, 7.5 mg/kg) to prevent motion artifacts. It has been shown that barbiturates do not have any influence on the regional distribution of CBF or on default mode resting state network (Werner 1995; Zilbovicius et al. 2000; Mishra 2002; Fransson et al. 2007; Doria et al. 2010).

### Data processing and treatment

MRI images were pre-processed using Statistical Parametric Mapping (SPM8 software, Welcome Department of Cognitive Neurology London www.fil.ion.ucl.ac.uk/spm/software/spm8) implemented in Matlab (Mathworks Inc., Sherborn, MA, USA) and analyzed using a voxel-based approach (See SI Appendix, SI Methods for details). Native 3D-T1-weighted images were segmented into gray matter, white matter and cerebrospinal fluid using the Infant Brain Probability Templates (https://irc.cchmc.org/software/infant.php). The unified segmentation enables spatial normalization, tissue segmentation and bias correction within the same generative model (Ashburner and Friston 2005). A postprocessing visual quality control was performed by two independent investigators (PA and JMT) on the grey matter maps to ensure the quality of the segmentation. The ASL images were co-registered to the corresponding native gray matter images using a normalized mutual information cost function (separation: 4 and 2 mm, tolerance: from 0.02 to 0.001, histogram smoothing: 7 x 7 mm). After visual inspection of the co-registration, the ASL images were then spatially normalized using the deformation matrices from the segmentation process. The resulting ASL images were smoothed using an isotropic Gaussian filter of 10 mm. ASL acquisition provides a high-quality image of quantitative CBF. Motion in ASL acquisition is mainly characterized by signal outside of the brain, often recognizable as signal from layers of skin or fat, that can be detected by on-the-fly expert visual analysis. Therefore, we performed a two steps quality control. The first one by an expert radiologist right after acquisition (NB) and the second one by an imaging processing expert engineer (HL) before pre-processing to discard images with artifacts such as motion, aliasing, ghosting, spikes, low signal to noise ratio.

### Image analysis

We normalized rest CBF within the ASL images by the mean CBF measured within the basal ganglia to avoid major variations in rest CBF due to cardiac blood flow (Licht et al. 2004; Varela et al. 2012) and blood pressure labilities (Hardy et al. 1997). The basal ganglia was specifically chosen in our study as it is one of the earliest structures to matures (Chugani and Phelps 1986) and regression analyses did not show any age-related variations in the rest CBF of this region within our age range (beta= 5.2 x 10^−4^ unit/day, t_(50)_ = 0.045, p = 0.96). The regional rest CBF was expressed as percentage of basal ganglia rest CBF and presented in arbitrary unit.

We then performed whole-brain voxel-wise analyses of the 52 images within the general linear model framework using SPM8. Age was entered as covariate in a multiple regression model. The analyses were constrained to gray matter tissue only by thresholding the analysis mask to 40% of the mean gray matter image of our sample.

We also extracted mean rest CBF from 92 regions of interest (hemispheres and regions) using the AAL parcellation toolbox (Tzourio-Mazoyer et al. 2002). We matched the AAL parcellation to our sample by spatially normalizing the MNI single subject MRI brain to the Infant Brain Probability Templates. In addition, a further analysis was performed by selecting and merging regions of interested based on their relevance in term of development. We selected the hippocampus, the amygdala, the thalamus, the primary visual and auditory cortices, the insula, the superior temporal cortex. We formed the sensorimotor cortex by merging the precentral and postcentral regions, and the prefrontal cortex by merging the inferior, middle and superior frontal regions and the gyrus rectus. All analyses were performed using R version 3.6.1 (http://cran.r-project.org) and ggplot2 (3.2.1) and lme4 (1.1-21) packages. Age-related regressions were assessed using linear mixed models to account for the intra-subject left and right hemisphere measurements. Age, hemisphere and age-by-hemisphere interaction were entered as fixed effects and subject as nested random effect. We corrected for multiple comparisons using Bonferroni correction by multiplying the p values by the number of regions. Finally, we performed inter-regional correlation analyses of the rest CBF values to derive a Pearson’s correlation coefficient for each pair of regions and to build an inter-regional correlation matrix. The ordering of the correlation matrix was based on correlations with the first principal component of the same matrix.

## Results

The relative values of global rest CBF increased with age from 3 to 12 months in the right (b = 0.0010 unit/day, t_(55.71)_ = 6.64, p = 1.36E-08) and in the left hemisphere (b = 0.00078 unit/day, t_(55.71)_ = 5.34, p = 1.74E-06) with a greater age-related increase in the right as compared to the left (p = 0,0074, see Figure 1 and Table 1).

**Figure 1:**
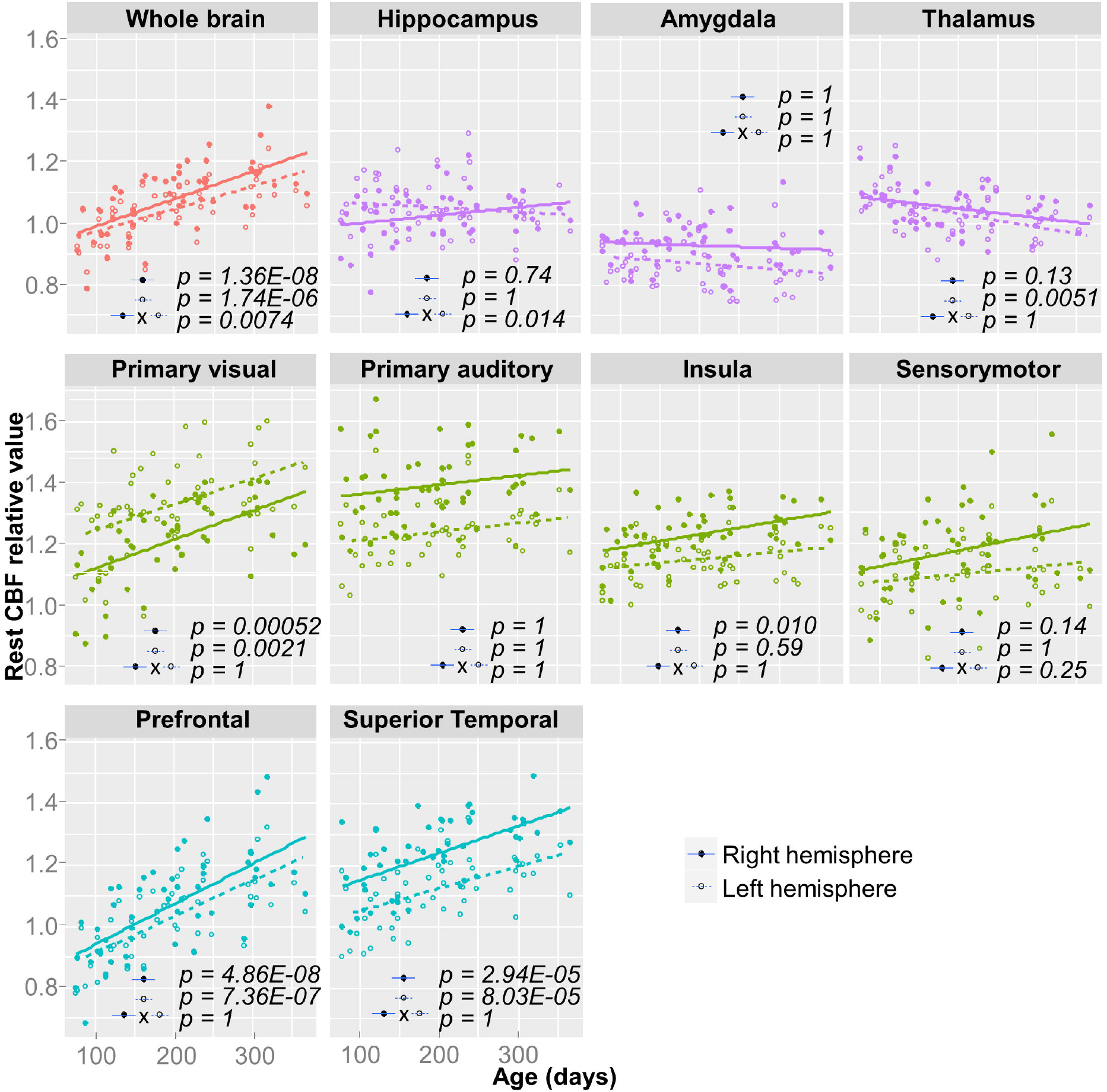
Age-related changes of the rest CBF values in predefined regions of interest between 3 and 12 months of age. The whole brain in red, subset of stable subcortical regions (hippocampus, amygdala and thalamus) in purple, subset of early maturing cortical regions (primary visual and auditory cortices, insula and sensorimotor cortex) in green, subset of late maturing cortical regions (prefrontal and superior temporal cortices) in blue. Each dot represents a subject, and each line represents the estimated regression based on a linear model for the left (empty dots and dashed line) and right (filled dots and solid line) hemispheres. The rest CBF values are normalized by the rest CBF measured within the basal ganglia and presented in arbitrary unit.

**Table 1:**
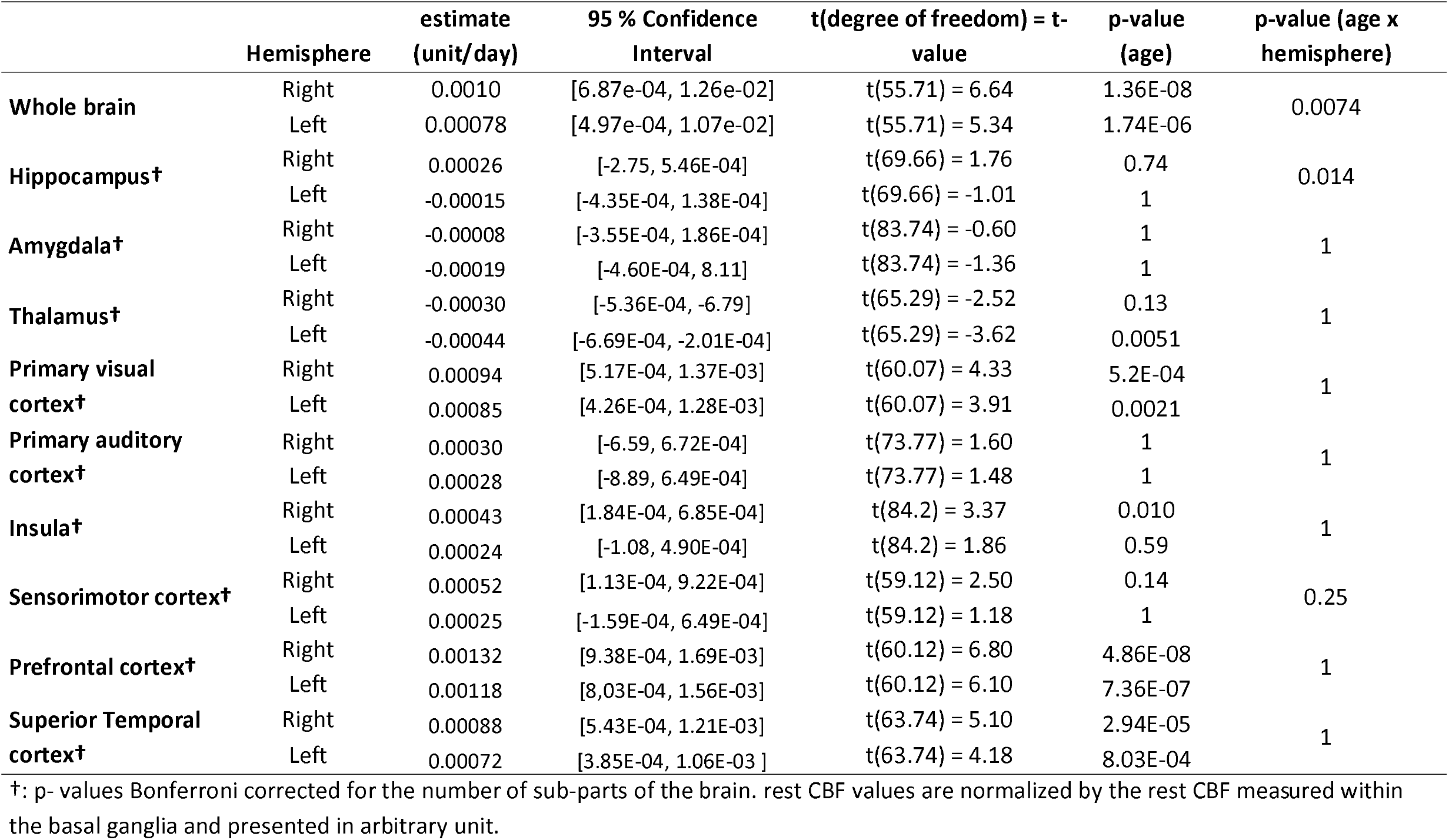
Age-related changes of the rest CBF values between 3 and 12 months of age.

Qualitative analysis of the whole-brain voxel-wise maps showed a regionally heterogeneous age-related increase of the relative rest CBF values (see Figure 2 and SI Appendix, Movies S1 to S4). The highest rest CBF at 3 months were observed within the sensorimotor and the primary visual cortices. The age-related increase in rest CBF progressed spatially from these regions. From the calcarine fissure, the rest CBF increased toward the visual associative regions up to the supramarginal and the precuneus regions. From the primary motor and sensory cortices, the rest CBF increased toward both the anterior and the posterior part of the brain. Anteriorly, through the anterior cingulate and the prefrontal cortices; posteriorly, through the insula and the superior temporal cortices. In contrast, the rest CBF was stable within the thalamus, the amygdala and the hippocampus. Between 9 and 12 months, the rest CBF increase was predominantly seen in the temporal and the prefrontal cortices. The regional right over left rest CBF asymmetry remained present throughout the whole studied period.

**Figure 2:**
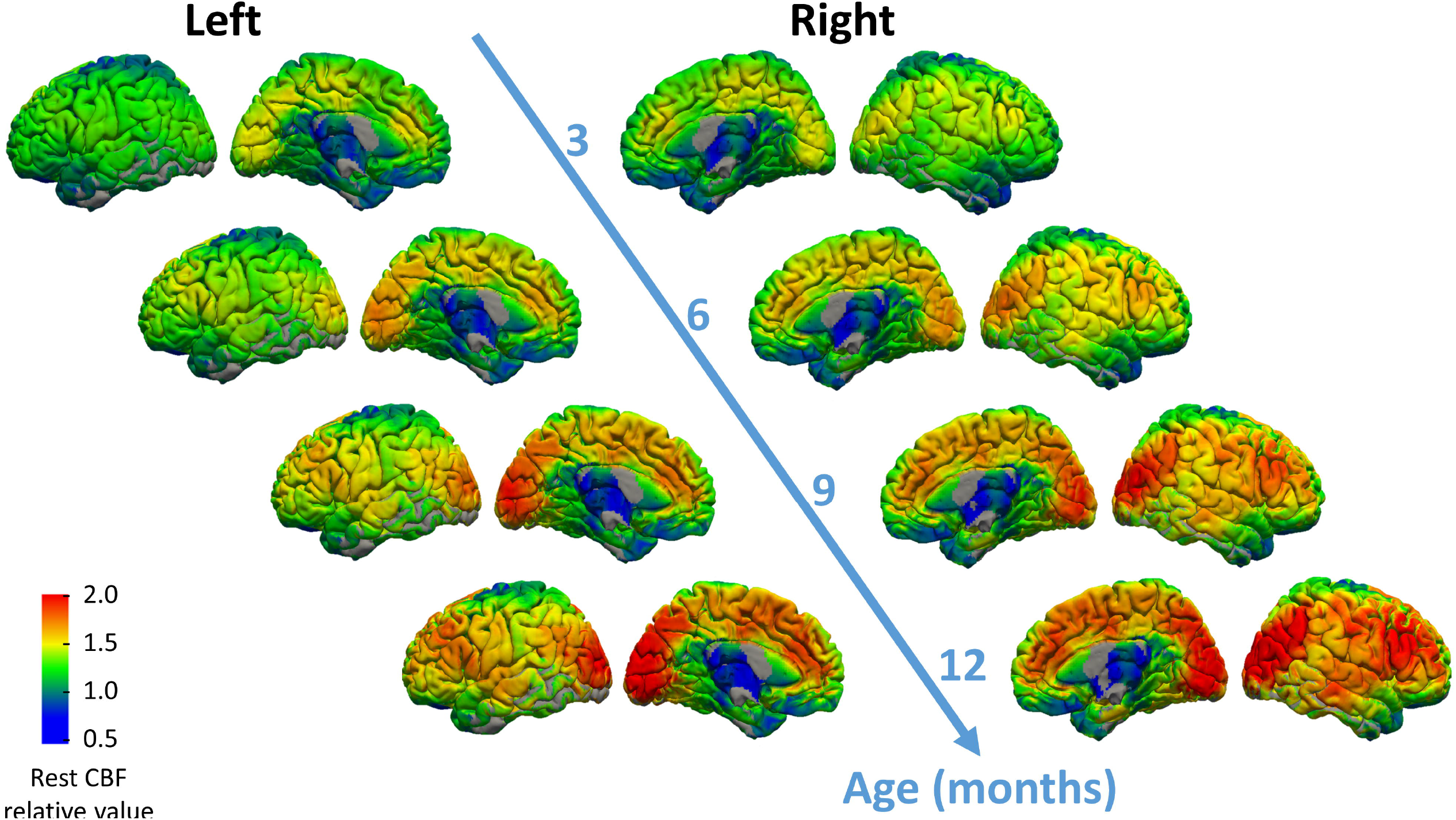
rest CBF values at 3, 6, 9 and 12 months of age displayed on the medial and lateral view of the left and right hemispheres. The rest CBF values are normalized by the rest CBF measured within the basal ganglia and presented in arbitrary unit. Surface rendering was done using mri_vol2surf from freesurfer (https://surfer.nmr.mgh.harvard.edu/).

Quantitative analysis within the predefined regions of interest showed different trajectories of local rest functional maturation (see Table 1 and Figure 1). First, in a subset of subcortical regions including the hippocampus (right: b = 0.00026 unit/day, t_(69.66)_ = 1.76, p = 0.74; left: b = -0.00015 unit/day, t_(69.66)_ = -1.01, p = 1), the amygdala (right: b = -0.00008 unit/day, t_(83.74)_ = -0.60, p = 1; left: b = -0.00019 unit/day, t_(83.74)_ = -1.36, p = 1) and the thalamus (right: b = -0.00030 unit/day, t_(65.29)_ = -2.52, p = 0,13; left: b = -0.00044 unit/day, t_(65.29)_ = -3.62, p = 0.0051), the age-related rest CBF maturation through the first year of life remained stable indicating already matured regions at 3 months old. Second, the subset of cortical regions including the primary visual (right: b = 0.00094 unit/day, t_(60.07)_ = 4.33, p = 5.2E-04; left: b = 0.00028 unit/day, t_(60.07)_ = 3.91, p = 0.0021) and primary auditory cortices (right: b = 0.00030 unit/day, t_(73.77)_ = 1.60, p = 0.94; left: b = 0.00028 unit/day, t_(73.77)_ = 1.48, p = 1), the insula (right: b = 0.00043 unit/day, t_(84.2)_ = 3.37, p = 0.010, left: b = 0.00024 unit/day, t_(84.2)_ = 1.86, p = 0.59) and the sensorimotor cortex (right: b = 0.00052 unit/day, t_(59.12)_ = 2.50, p = 0.14; left: b = 0.00025 unit/day, t_(59.12)_ = 1.18, p = 1) presented a small age-related rest CBF increase indicating early maturational process. Third, a subset of cortical regions including the prefrontal (right: b = 0.00132 unit/day, t_(60.12)_ = 6.80, p = 4.86E-08; left: b = 0.00118 unit/day, t_(60.12)_ = 6.10, p = 7.36E-07) and the superior temporal cortices (right: b = 0.00088 unit/day, t_(63.74)_ = 5.10, p = 2.94E-05; left: b = 0.00072 unit/day, t_(63.74)_ = 4.18, p = 8.03E-04) presented a high age-related rest CBF increase indicating late maturational process. Faster right over left age-related rest CBF increase was more pronounced within the hippocampus (p = 0.014). Finally, the rest CBF inter-regional correlation matrix between the predefined regions showed a cluster of highly correlated regions with the following rank order: Superior temporal, prefrontal, insula, sensorimotor and primary auditory cortices (see Figure 3).

**Figure 3:**
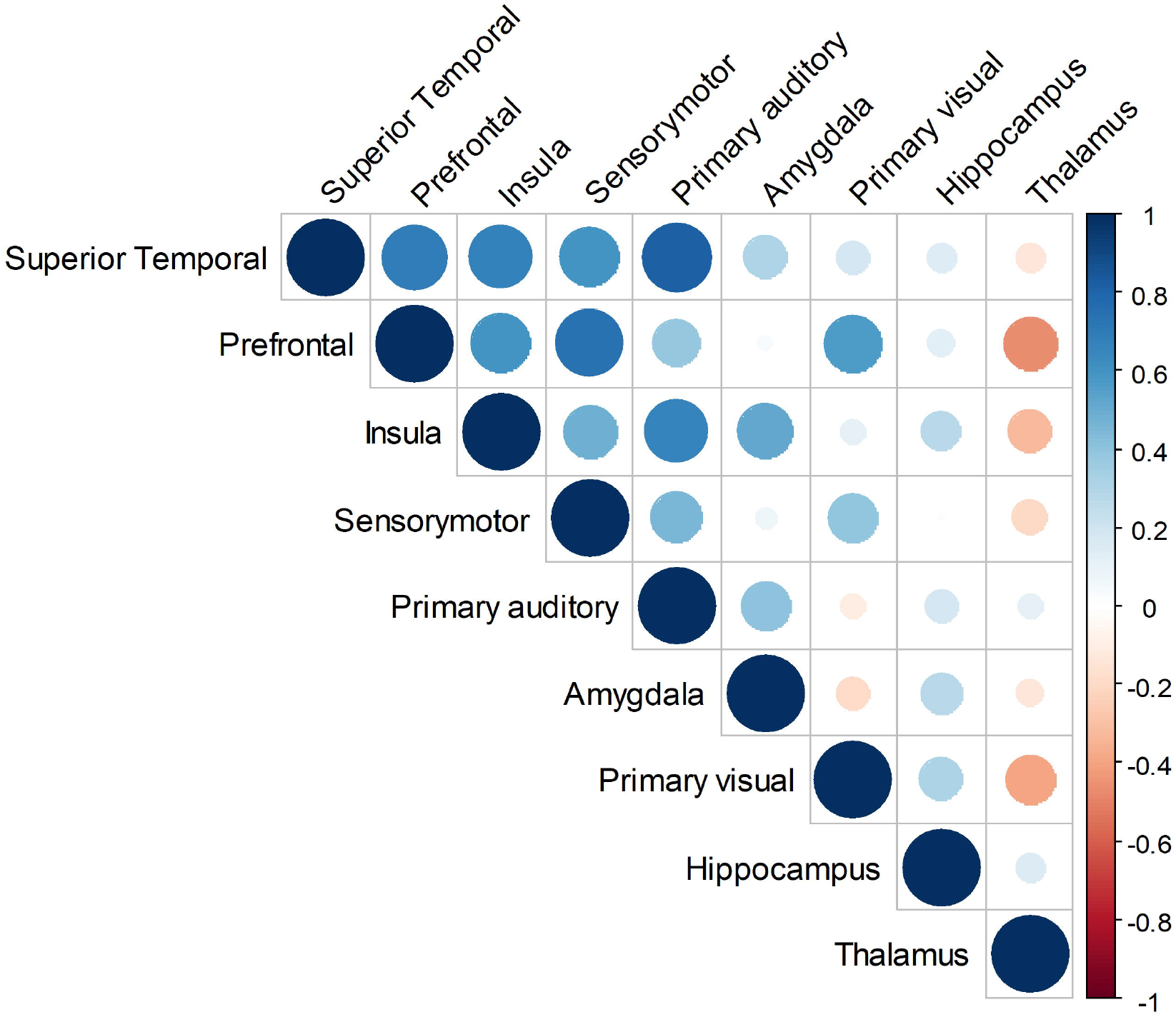
Correlogram of the correlation matrix for the rest CBF values in the predefined regions of interest. Size and color of the circle represent the Pearson’s correlation coefficients. Correlation ordering is based on correlations with the first principal components of the same matrix (i.e. similarity measure).

The age-related rest CBF changes computed for the exhaustive list of 45 regions of interest are available in SI Appendix, Table S3.

## Discussion

Our study shows for the first time the dynamics of local rest functional brain maturation throughout the first year of life using a non-invasive imaging method. Global rest CBF increased significantly from 3 to 12 months of age and this increase was more pronounced in the right than in the left hemisphere. Qualitative and quantitative analyses revealed marked regional differences in local functional brain maturation. Subcortical structures such as basal ganglia, thalamus, amygdala and hippocampus cortices are stable at 3 months. At the cortical level, we observed two different maturational trajectories: first, a set of regions with a low age-related rest CBF increase between 3 to 12 months, including the primary auditory/visual cortices, the sensorimotor cortex and the insula. A second set of regions a high age-related increase rest CBF increase between 3 to 12 months, including the superior temporal and prefrontal cortices.

The increase in global rest CBF from 3 to 12 months of age that we describe here is consistent with pioneers PET and SPECT studies showing increase in rest metabolism and CBF during the same period (Chugani et al. 1987; Chiron et al. 1992). Furthermore, we highlighted a hemispheric functional maturational asymmetry, with greater right than left global rest CBF increase during the first year. This agrees with previous studies that showed greater right than left rest CBF for these regions at birth (Lin et al. 2013) and from 1 year to 3 years old (Chiron et al. 1997), supporting the hypothesis that the right hemisphere functionally matures earlier than the left.

Globally, our findings are in accordance with results from prior research based on histology, structural and rest functional brain imaging that has revealed distinct maturation trajectories of cortical regions and brain networks over the first year of life (Chugani and Phelps 1986; Gilmore et al. 2012; Smyser and Neil 2015; Zhang et al. 2019). Firstly, at the histological level, post-mortem data showed that the time course of synaptogenesis differs across cortical regions. Indeed, a burst of synapse formation occurs between 3 and 4 months within primary visual, auditory cortices somatosensory cortices, which appeared already mature at 3 months of life (Marin-Padilla 1970; Huttenlocher 1979, 1990; Michel and Garey 1984). In non-human primates, Rakic and colleagues have shown a synchronic synaptogenesis in the visual, somatosensory, motor, and prefrontal areas (Rakic et al. 1986). In humans, synaptogenesis in the prefrontal cortex begins about the same time as in visual cortex, but it does not reach its peak period until age 8 months, continuing thereafter through the second year of life (Kostovic et al. 1995; Huttenlocher and Dabholkar 1997). Using quantitative electron microscopy in non-human primates, this remarkable overproduction followed by elimination of synapses in the prefrontal cortex have also been described (Bourgeois et al. 1994). These congruent findings strengthen our results as synaptic density is coupled to rest CBF. Secondly, concerning myelination, microstructural MRI maturational studies described a global maturation pattern characterized by early maturation of the sensorimotor cortex, followed by the other sensory cortices and then the associative cortices, including the prefrontal cortex (Dubois et al. 2008; Deoni et al. 2011). Finally, recent data obtained with resting-state functional MRI studies allowed to describe maturational changes of functional networks during the first year of life (Smyser et al. 2010; Gao et al. 2015; Smyser and Neil 2015; Wen et al. 2019). Especially, Gao et al. have described a maturation sequence starting with primary sensorimotor/auditory and visual then attention/default-mode, and finally executive control, prefrontal, networks (Gao et al. 2015). These different sequences of functional network maturation fit with and complement our results. Therefore, data coming from our study and previous rs-fMRI studies contribute to map a timetable of functional brain maturation during the first year of life.

Importantly, the spatial resolution of the ASL images allowed an accurate mapping of the age-related rest CBF changes. Consequently, we were able to describe insular local functional maturational evolution, which reaches its one-year pattern rather early, during the first months of life. A well-established literature and recently anatomical and resting-state functional MRI studies describe early human cortical development in areas close to the insula and radiating outward (Alcauter et al. 2015). This early insular maturation fits with its role in the integration of interoceptive stimuli, such as coolness, warmth and distension of the bladder, stomach or rectum (Craig 2009), but also in the integration of external stimuli, notably pain (Mazzola et al. 2009). In addition, it is highly pertinent, since the insula is a key structure for the baby’s development and essential to baby’s survival.

The spatial resolution improvement also allowed us to describe for the first time a remarkably synchronous increase in rest CBF between the prefrontal and superior temporal cortices (see Figure 3), both main components of the called “social brain” (Brothers et al. 1990). The late maturation of the prefrontal cortex had been previously described by structural and functional brain imaging studies (Chugani and Phelps 1986; Gilmore et al. 2012). Noticeably, we describe here a late maturation within the posterior temporal regions during the first year of life, particularly within the posterior superior temporal sulcus, a region known to be highly implicated in social cognition (Zilbovicius et al. 2006). Interestingly, the late and synchronous maturation of these two cortical structures corresponds to the remarkable development of the baby’s social skills through the first year of life.

To the best of our knowledge, only 2 studies using ASL imaging have focused on brain development during the first year of life, but a comparison with our results is limited due to important differences in their methodological approaches. Duncan et al. studied a sample of 61 infants within a very narrow age-range from 3 to 5 months (Duncan et al. 2014). Their main results describing a significantly greater rest CBF in the sensorimotor and occipital regions compared with the dorsolateral prefrontal in this age-range are in accordance with our results. In the second study, combining region of interest (ROI) and whole-brain analyses on rest CBF, Wang et al. investigated a group of 8 7-month-old infants to a group of 8 13-month-old infants (Wang et al. 2008). Although they showed rest CBF increase in the 13-month-old group compared to the 7-month-old group mainly located in the frontal lobe, they did not examine directly the age-related rest CBF slopes.

This study has some limitations. First, we used a linear model for data analysis. Although cubic and quadratic fitting models did not improve our statistical models, it is improbable that a linear model exactly fits functional cortical maturation. This issue can be addressed in future studies by adding more and older subjects to further investigate the postnatal brain rest functional maturation trajectory. Second, due to their age, all infants received light premedication before the MRI to prevent motion artifacts, and all the scans were acquired during sleep. No significant influence neither on the regional distribution of CBF (Fransson et al. 2007; Doria et al. 2010; Carsin-Vu et al. 2018) nor in the default-mode network connectivity (Greicius et al. 2008) has been reported to this premedication. Third, it is important to stress that CBF is an indirect surrogate marker of focal neural activity by providing two important metabolic substrate, oxygen and glucose, that are critically important to neurons, synapses and astrocytes. Although, this neuro-hemodynamic coupling plays an important role in brain development through dendritic sprouting, axonal growth, synapse formation and vascular patterning, it does not necessarily imply a maturity of the resulting neural activity. Finally, our study was performed in a clinical pediatric population. To ensure that it could be comparable with a non-clinical population, we discarded all clinical indications for MRI that could affect brain anatomy, function and further neurodevelopmental disorders. In addition, all scans were strictly normal, and follow-up confirmed a normal psychomotor development.

Defining typical trajectories of brain maturation provides references for a better understanding of neurodevelopmental disorders and preterm effects on further brain maturation. Because CBF reflects regional changes in synaptic density, ASL offers a noninvasive approach to studying local brain function. Furthermore, the recent possibility to implement ASL perfusion-based functional connectivity in conjunction to regular resting state BOLD connectivity should be investigated for characterizing spatiotemporal and quantitative properties of cerebral networks during brain maturation (Chen et al. 2015). In conclusion, to our knowledge, our study is the first to describe and characterize dynamics local functional brain maturation during the first year of life and provide insight into an important and vulnerable neurodevelopmental period.

## Notes

The authors declare no competing interests. The data that support the findings of this study are available on request from the corresponding author. The data are not publicly available due to them containing information that could compromise research participant consent.

## Corresponding author

Hervé Lemaître, PhD, Groupe d’Imagerie Neurofonctionnelle, Institut des Maladies Neurodégénératives (CNRS UMR 5293), Université de bordeaux, Centre Broca Nouvelle-Aquitaine, 146 rue Léo Saignat - CS 61292 - Case 28, 33076 Bordeaux cedex, France

## Notes

### Competing Interest Statement

The authors have declared no competing interest.

