## Supplementary Information Text for "Rest functional brain maturation during the first year of life"

**SI Methods.**

**MRI acquisition.** All MRI exams included T1-weighted and ASL sequences and were acquired on a General Electric Signa 1.5T MRI scanner in the Necker-Enfants-malades hospital. We acquired a 3D pseudocontinuous arterial spin labelling (3D pcASL) sequence using a fast spin echo acquisition with spiral filling of the K space (TR/TE: 4453/10.96 msec, 8 spiral arms x 512 sampling points, post-labelling delay: 1025 msec, flip angle: 155°, matrix size: 128 x 128, slice thickness: 4 mm, Field of View: 24 x 24 cm, 40 contiguous axial slices, duration: ≈ 5 mins, see SI Appendix, Table S2). CBF quantification from ASL tagged and control images was performed and warranted by General Electric 3D ASL software (Zaharchuk et al. 2009). Anatomical sequences were used only for spatial normalization of ASL sequences.

Zaharchuk G, Bammer R, Straka M, Shankaranarayan A, Alsop DC, Fischbein NJ, Atlas SW, Moseley ME. 2009. Arterial spin-label imaging in patients with normal bolus perfusion-weighted MR imaging findings: pilot identification of the borderzone sign. Radiology. 252:797–807.

Table S1. Clinical MRI indication by age group.

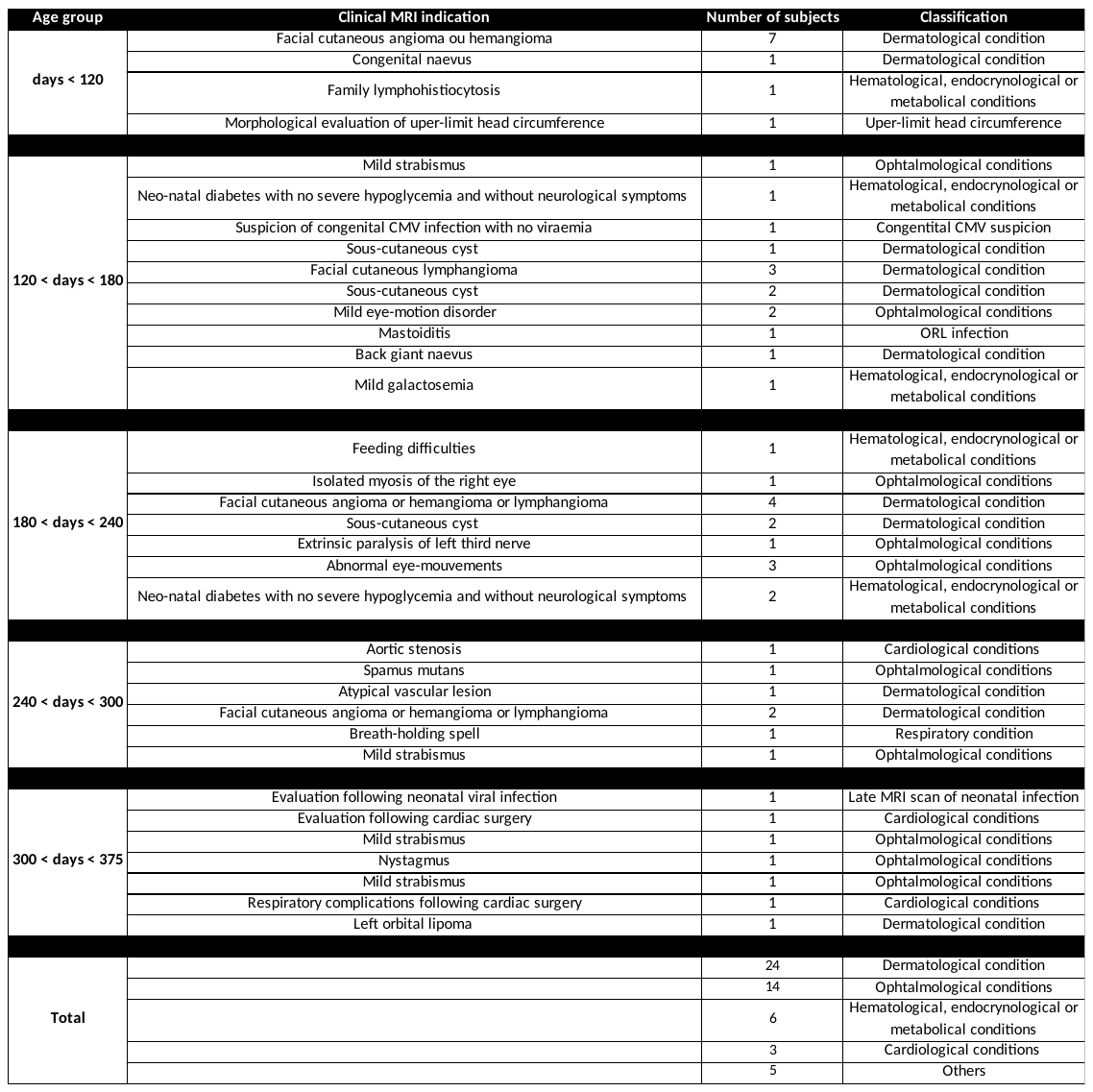

Table S2. Imaging parameters for the 3D pseudocontinuous ASL sequence.

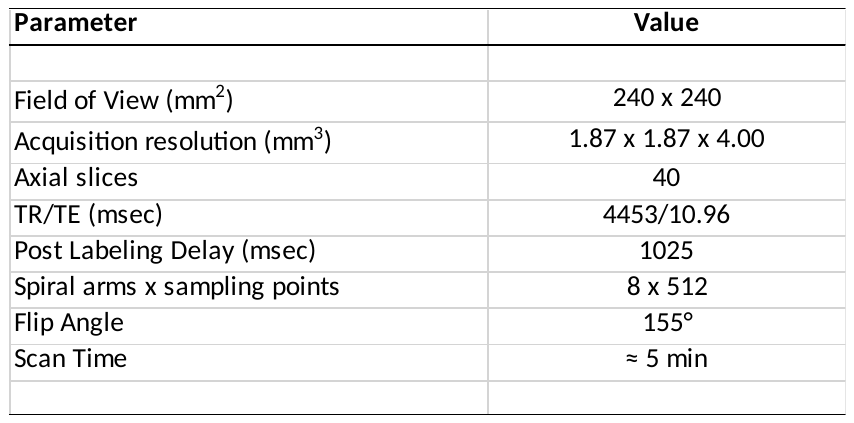

Table S3. Age-related changes of the rest CBF values between 3 and 12 months of age for all regions.

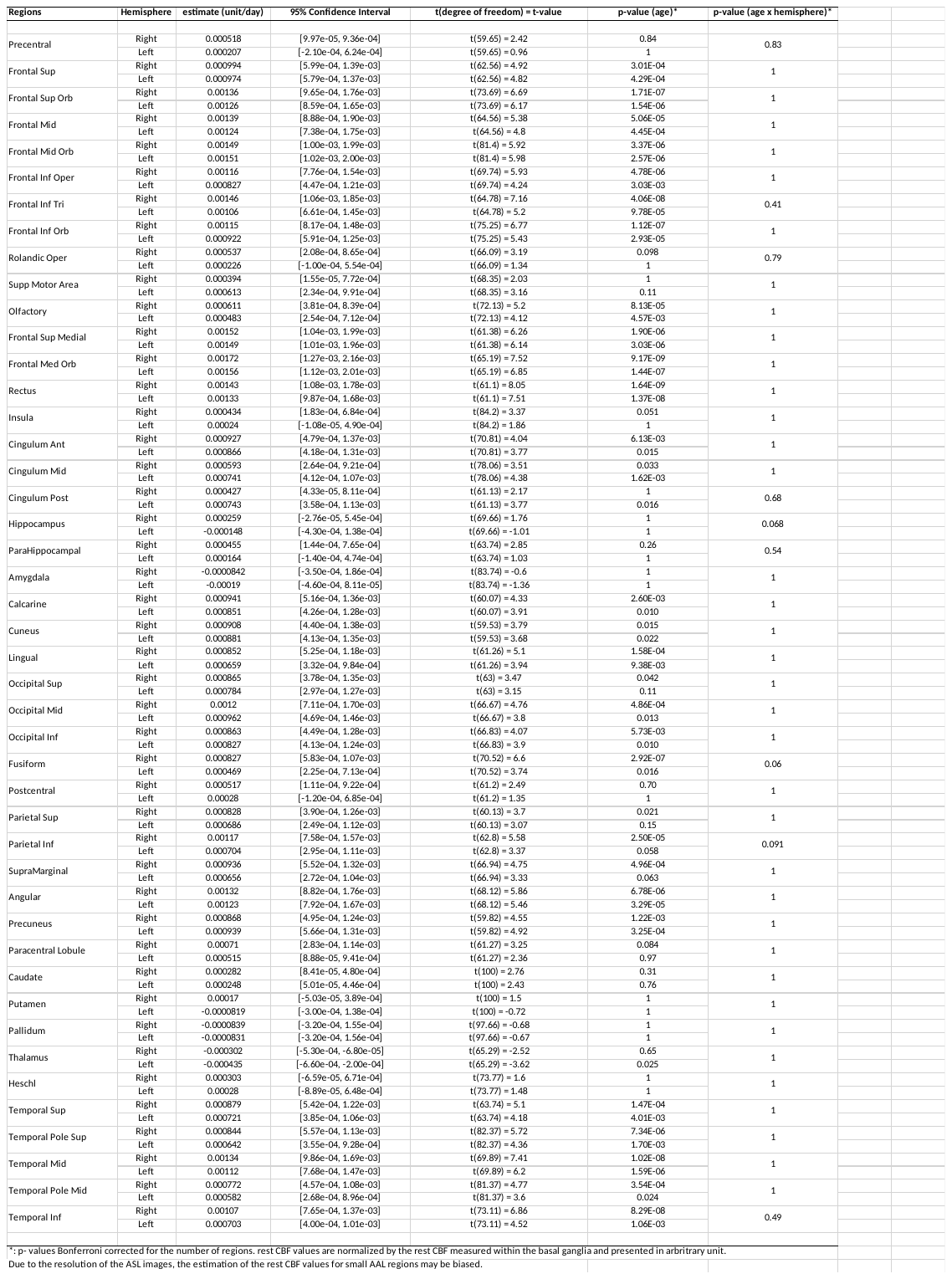

Movie S1. Video of variations in rest CBF between 3 and 12 months of age within the lateral view of the left hemisphere. The values observed correspond to the normalization of rest CBF values to rest CBF measured within the basal ganglia and presented in arbitrary unit.

Movie S2. Video of variations in rest CBF between 3 and 12 months of age within the lateral view of the right hemisphere. The values observed correspond to the normalization of rest CBF values to rest CBF measured within the basal ganglia and presented in arbitrary unit.

Movie S3. Video of variations in rest CBF between 3 and 12 months of age within the medial view of the left hemisphere. The values observed correspond to the normalization of rest CBF values to rest CBF measured within the basal ganglia and presented in arbitrary unit.

Movie S4. Video of variations in rest CBF between 3 and 12 months of age within the medial view of the right hemisphere. The values observed correspond to the normalization of rest CBF values to rest CBF measured within the basal ganglia and presented in arbitrary unit.
